# BBB damage in aging causes brain iron deposits via astrocyte-neuron crosstalk and Hepc/Fpn1 pathway

**DOI:** 10.1101/2021.07.01.450665

**Authors:** Mariarosa Mezzanotte, Giorgia Ammirata, Marina Boido, Serena Stanga, Antonella Roetto

## Abstract

During aging, iron accumulates in brain’s regions vulnerable to neurodegeneration: the cerebral cortex and the hippocampus. However, the mechanism of iron regulation in the brain remains scarce. Here, we demonstrated for the first time the involvement of the Hepcidin/Ferroportin1 pathway in brain iron metabolism during aging.

We demonstrated the alteration of BBB integrity, that leads to increased iron permeability and deregulation of iron homeostasis during aging. We found that brain iron overload drives Hepcidin upregulation and, consequently, the inhibition of the iron exporter Ferroportin1, neuroinflammation and oxidative stress. Moreover, both in the cerebral cortex and hippocampus Ferroportin1 colocalizes with astrocytes, while the iron storage protein ferritin light-chain with neurons. This differential distribution suggests that astrocytes mediate iron shuttling and neurons are unable to metabolize it. Furthermore, we observed NCOA4-dependent ferritinophagy of ferritin heavy-chain isoforms determining the increase of light-chain enriched ferritin heteropolymers that are more efficient as iron chelators. Altogether, these data highlight the involvement of the Hepcidin/Ferroportin1 axis and NCOA4 during mice aging as a response to a higher iron influx to the brain.

## Introduction

Iron is an essential metal in many cellular and biological processes: oxygen transport, oxidation-reduction reactions, ATP production, DNA biosynthesis and repair. Conversely, it can generate Reactive Oxidative Species (ROS) by Fenton reaction, contributing in this way to the pathophysiology of many diseases characterized by iron overload (Ganz 2012).

Cells need a balance between iron availability for several biochemical processes, and iron deposits. This homeostatic situation is guaranteed by the action of different proteins involved in iron transport into the cells: Transferrin (Tf), Transferrin Receptors (TfR1) and Divalent Metal Transporter 1 (DMT1); iron export, through the action of the main cellular iron exporter Ferroportin 1 (Fpn1) (Ginzburg 2019) and storage, carried out by cytosolic ferritin (Ft) heteropolymer, composed of 24 subunits of ferritin heavy (Ft-H) and light (Ft-L) chains that can store up to 4500 iron atoms (Arosio 2009; Morello 2009).

Nonetheless, iron trafficking is a dynamic process where Tf plays a major role in iron relocation between sites of absorption, recycling and storage and sites of utilization. Transferrin-iron (Tf-Fe2) complex binds to Transferrin Receptor 1 (TfR1), which is ubiquitously expressed in all cells.

However, the highest concentration of TfR1 is detected in erythroid precursors since hemoglobin producing cells request the highest amount of iron in the body (Ginzburg 2019).

Iron content and availability in the body are regulated by Hepcidin (Hepc), a peptide produced mainly by hepatocytes. Hepc interacts with the cellular iron exporter Fpn1, causing its degradation, and iron retention inside the cells (Sangkhae 2017). Consequently, Hepc decreases the amount of iron in the serum (Ganz 2013), it controls intestinal iron uptake and its release from splenic macrophages according to the body need (Sangkhae 2017). In iron deficiency conditions (i.e. anemia, hypoxia, ineffective erythropoiesis) Hepc level decreases; on the contrary, its expression increases when the content of iron in the organism is sufficient or during inflammatory processes (Sangkhae 2017; Roetto 2018).

In the brain, iron is essential for different metabolic processes and it regulates important functions such as neurotransmission, myelination and division of neuronal cells (Moos 2007). Iron enters into the brain crossing the Blood Brain Barrier (BBB), which is the pivotal protective filter to avoid inappropriate levels of iron in the brain (Mills 2010; Thirupathi 2019). Iron shuffling into the brain is mediated by the same transporters that are acting in the peripheral tissues; iron up-take is mediated by TfR1 that is expressed on the luminal side of brain capillaries (Bien-Ly 2014; McCarthy and Kosman 2014) where it binds circulating Tf-Fe2 to promote iron uptake into brain microvascular endothelial cells (BMVECs) through coated pits endocytosis (Mills 2010). Iron is released into the cytoplasmic space and exported through the abluminal membrane by unknown mechanisms in which Ferroportin1 may be involved (Donovan 2000) as well as other transporters (Ward 2010).

Several other genes that regulate iron homeostasis are expressed in the murine Central Nervous System (CNS), including Iron Regulatory Proteins (IRPs) (Leibold 2001), Ft (Moos 1996), neogenin-1 (NEO-1) (Rodriguez 2007). Indeed, iron enters into neurons, microglial and cells of the choroid plexus bound to TfR1. It has been shown that Hepc is also present in the brain, both in mature astrocytes and oligodendrocytes (Vela 2018), where it plays a role in the control of iron amount together with its own iron regulatory proteins (Ward 2010). It is not yet clear whether all the Hepc acting on Fpn1 in the brain could have been produced in the liver (Vela 2018). Although the peptide size and its amphipathic cationic chemical structure (Bulet 2004) would allow hepatic Hepc to pass the BBB, it has been shown that there is an endogenous cerebral Hepc expression (Zechel 2006) and that it responds to brain iron (BI) state (Pellegrino 2016).

Moreover, in vivo studies showed that synthetic Hepc injection in rats’ brains caused a decrease in Fpn1 levels, demonstrating that Hepc/Fpn1 axis works also in the brain (Raha-Chowdhury 2015).

As in other organs, BI balance should be carefully maintained in order to avoid neurotoxicity induced by excessive concentrations of free iron (Ward 2010). It was shown that an increase in BI levels could be due to several conditions such as inflammation, iron redistribution inside the brain, iron unbalance (Conde 2006; Farrall 2009) and, consequently, BBB permeability increases because of the release of inflammatory mediators, free radicals, vascular endothelial growth factor, matrix metalloproteinases and microRNAs (miRNAs) (Almutairi 2016). Interestingly, the processes of aging could also be responsible for misregulated iron metabolism (Ward 2010). Indeed, it is known that iron amount increases during aging and that several neurodegenerative disorders (NDs), such as Parkinson’s disease (PD), Alzheimer’s disease (AD) and Multiple Sclerosis (MS), are associated with inappropriate iron accumulation and oxidative stress that can quicken ferroptosis, an iron-dependent form of cell death (Rouault 2013; Santana-Codina 2018, Quiles Del Rey 2019). During the physiological process of aging, iron accumulation is found in different brain regions, known to be more vulnerable to age-dependent neurodegeneration (Mills 2010).

Different studies suggested that cerebral cortex (Ctx) iron increase would have a significant impact on cerebral functioning: indeed iron has been proposed as a marker of neurodegeneration (Biasiotto 2016; Buijs 2017).

Recently, a new protein named Nuclear Receptor Coactivator 4 (NCOA4) which is involved in iron metabolism has been identified: it is a cargo protein which promotes selective autophagic ferritin degradation (Mancias 2014). As a consequence of NCOA4 binding to Ft-H, ferritin is carried to the lysosome where it is degraded and iron is released in the cytoplasm, modulating in this way intracellular iron regulation, in a process termed “ferritinophagy” (Mancias 2015; Santana-Codina 2018). Interestingly, NCOA4 levels are regulated by intracellular iron status (Mancias 2015) and recent works showed that the amount of NCOA4 changes according to the interaction with another protein, HERC2, an E3 ubiquitin protein ligase (Mancias 2015; Quiles Del Rey 2019). Moreover, Bellelli et al. described an iron overload phenotype in a NCOA4 knockout mouse model with increased level of Tf saturation, serum Ft, liver Hepc and an increase of Ft deposits (Bellelli 2016). Recently, an extra-hepatic function of NCOA4 was demonstrated (Nai 2021).

However, up to now, no data are available on NCOA4 and Hepc/Fpn1 expression levels and their functions in the brain during aging and/or neurodegeneration.

In the present study we investigated Hepc, Fpn1 and NCOA4 expression in a murine model during aging to unravel their possible involvement in BI overload.

## Results

### Iron amount and distribution in WT mice brain during aging

To verify BI amount, we evaluated BIC at each experimental time point (Fig.1A). Actually, BIC gradually increases during aging (Fig.1B).

**Figure.1.**
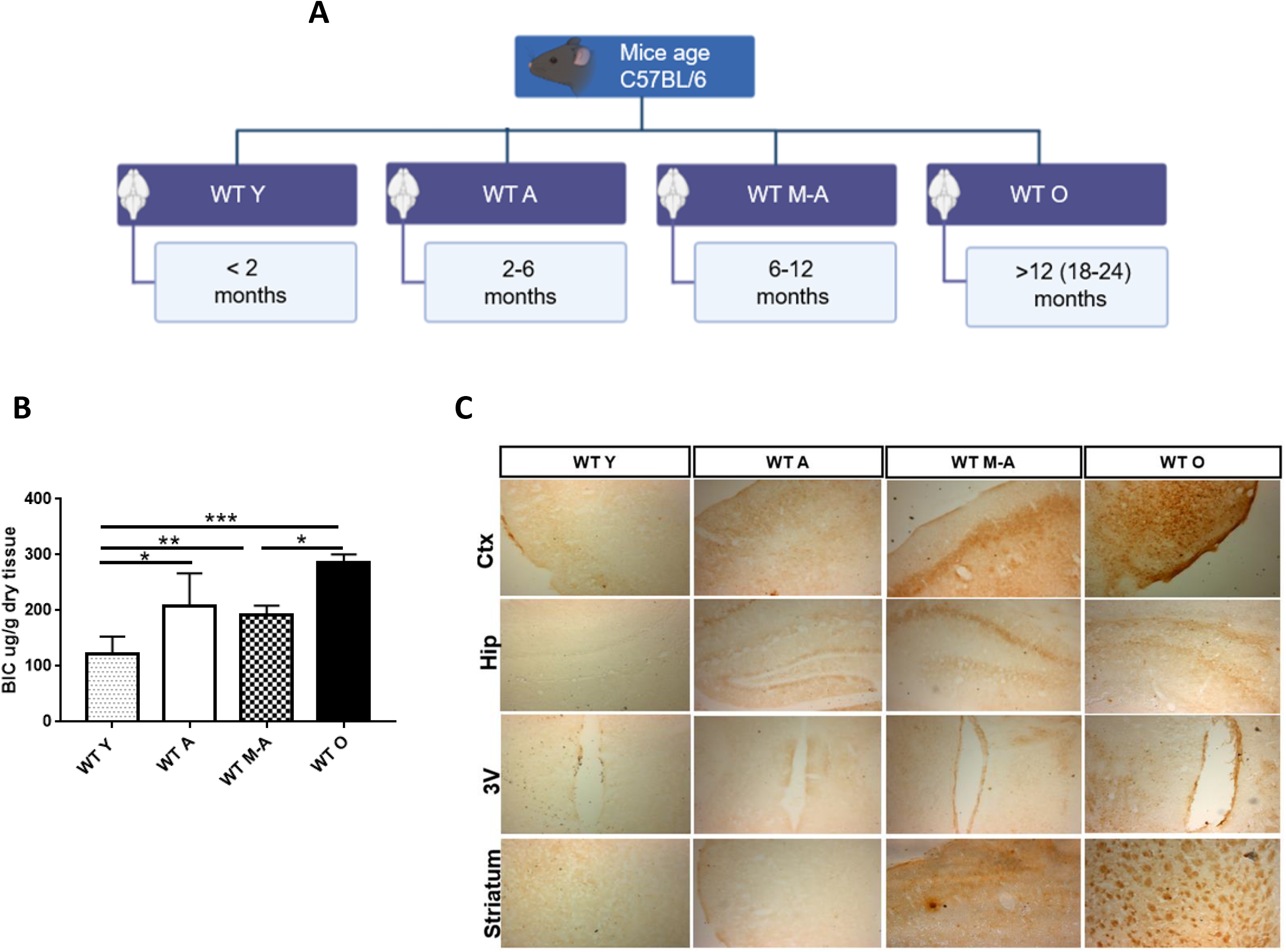
BI amount and accumulation in specific brain compartments during aging. (A) The Jackson Laboratory classification (https://www.jax.org): WT Y, Young; WT A, Adult; WT M-A, Middle-Aged; WT O, Old.; (B) Brain Iron Content (BIC) from mice total brain at different ages; (C) Sections of mice brain stained with DAB-enhanced Prussian Blue staining in cerebral cortex (Ctx), hippocampus (Hip), third ventricle (3v) and Striatum during ageing. Scale bars:10X; *statistically significant vs WT A control group *P <0.05; **P <0.01 ***P <0.001 using OneWay ANOVA followed by Bonferroni’s post hoc analysis or unpaired T-test.

In order to confirm and visualise the increase of iron in the brain and to investigate if iron deposition varies in different brain areas, we performed histochemical analysis with DAB-enhanced Prussian blue Perls’ staining. Indeed, the brain sections of WT O mice show an increased number of brown positive precipitates compared to WT A mice in specific parenchymal region such as Ctx, Hip CA regions, third ventricle (3V) and striatum (Fig.1C).

These results demonstrate that iron accumulates in the nervous tissue during physiological aging in specific areas of the brain.

### Iron-dependent inflammatory response and oxidative stress during aging

To investigate if the accumulation of iron in the brain led to a neuroinflammatory processes and if it could cause oxidative stress, we analysed the two main markers of these biological processes, SAA1 (Jang 2019) and Nrf2 (Jiang 2019) respectively.

SAA1 expression levels significantly become more than 20 times higher in WT O animals compared to WT A mice (Fig. 2A). Nrf2 expression levels also appear to constantly increase during aging (Fig. 2B).

**Figure.2.**
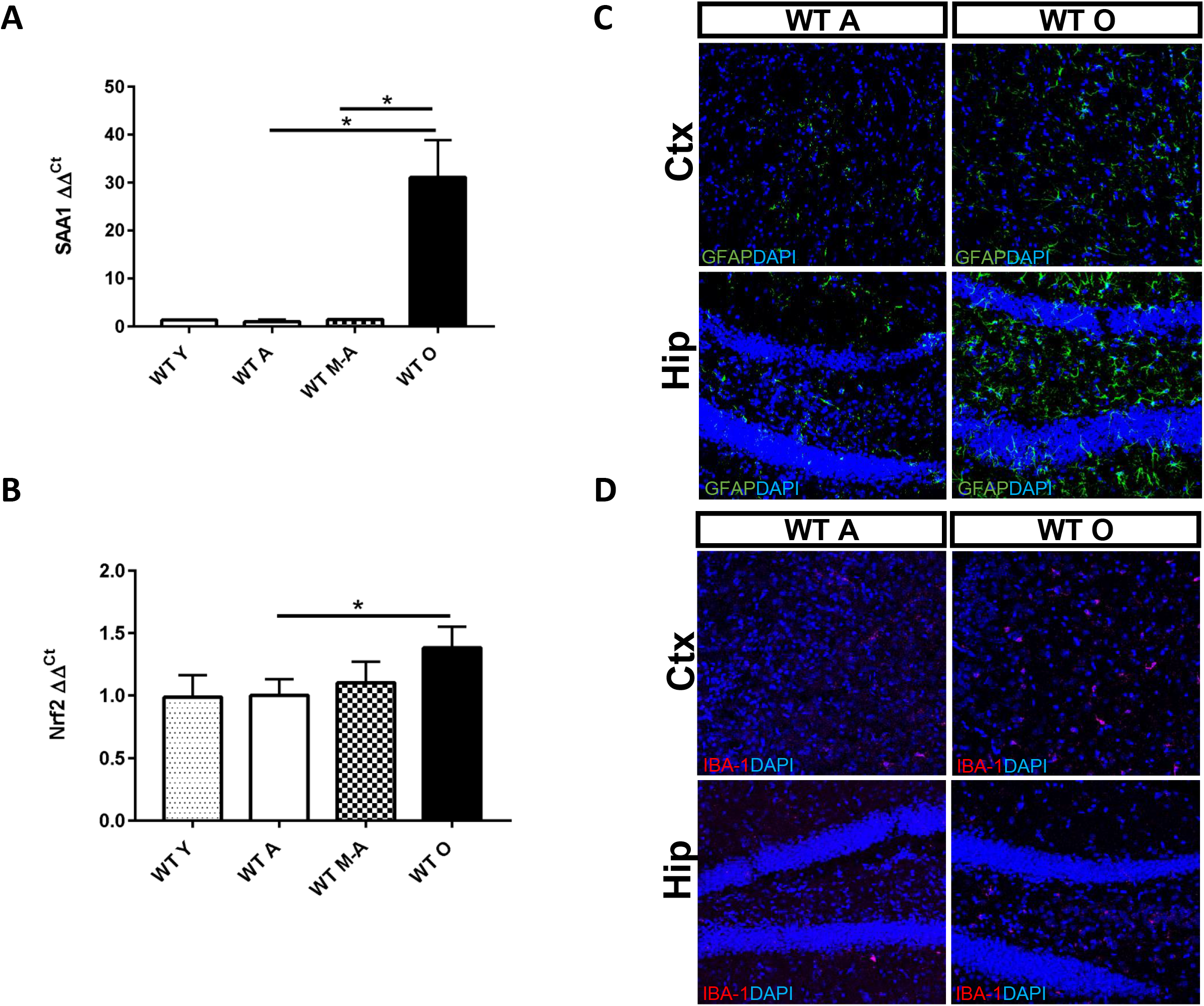
Inflammatory response activation and oxidative stress during aging. (A) Real-time PCR of Serum amyloid A1 (SAA1) in total brain from all genotypes. (B). Nuclear factor erythroid 2-related factor 2 (Nrf2) mRNA expression levels in total brain. The expression levels of SAA1 (A) and Nfr2 (B) were normalized to levels of β-glucuronidase (Gus-β) housekeeping gene (material and methods section); (C-D) Immunofluorescence anti-GFAP (green) and anti-IBA1 (pink) antibodies in cerebral cortex (Ctx) and hippocampus (Hip); 4,6diamidino-2-phenylindole (DAPI) (blue) was used to counterstain cell nuclei, scale bars:40X. *statistically significant vs WT A control group *P <0.05; **P <0.01 ***P <0.001 using OneWay ANOVA followed by Bonferroni’s post hoc analysis.

Moreover, since it is known that i) the expression of the intermediate filament protein (GFAP) is a morphological characteristic of astrocytes (O’Callaghan 2005) and ii) its activation is one common indicator of neuroinflammation in the CNS and it is involved in the progression of neurodegeneration in ischemia, Alzheimer’s disease (AD), Multiple Sclerosis (MS), Amyotrophic Lateral Sclerosis (ALS) and Parkinson’s disease (PD) (Mosley 2006; Glass 2010; Block and Hong 2005; Hirsch and Hunot 2009; Vezzani 2013), we performed immunohistochemistry to selectively label reactive GFAP-positive astrocytes. Moreover, we also checked the expression of IBA-1, a microglia/macrophage-specific calcium-binding protein which is also a key molecule in proinflammatory processes (Lier 2019). Indeed, we identified high astrocyte activation and an increased expression of microglia in both iron overloaded parenchymal regions of WT O mice, Ctx and Hip, compared to those of WT A mice (Fig. 2C and 2D).

These data show that the increase of iron in the brain triggers the neuroinflammatory and antioxidative stress response to iron transition from the systemic circulation to the brain.

### Hepcidin/Ferroportin1 pathway response to iron increase during aging

In order to evaluate the role of Hepc/Fpn1 axis and to verify if this pathway is also active during brain aging to promote iron retention into the cells, we measured both Hepc and Fpn1 levels in mice brain at different ages. We observed that Hepc gene expression significantly increases in WT M-A and WT O mice brain (Fig. 3A), while Fpn1, the target of Hepc, decreases (Fig. 3B). Due to the iron selective localization in the Ctx and Hip (Fig. 1C), we decided to confirm Fpn1 decrease in the same brain compartments. On the contrary of what happened in total brain, we found that Fpn1 increases during aging in both Ctx and Hip. In particular, immunofluorescence experiments demonstrated a clear increase of the protein in WT M-A and WT O mice Ctx and Hip compared to the same WT A mice areas (Fig. 3C).

**Figure.3.**
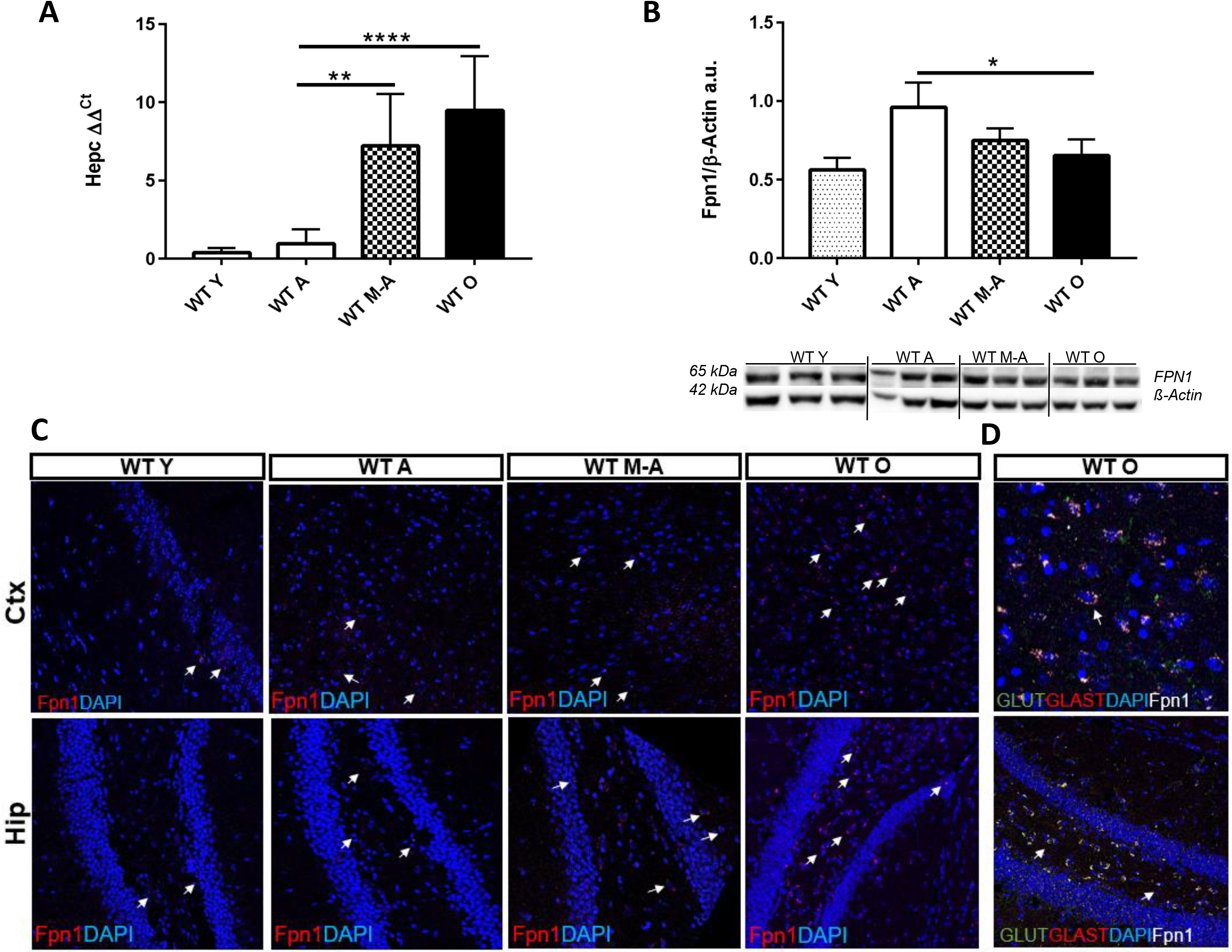
Hepc/Fpn1 pathway activation during aging. (A) Hepcidin (Hepc) transcription pattern, (B). Western blotting analysis and quantification of Ferroportin (Fpn1) in mice total brain during aging. (B) Immunofluorescence anti-Fpn1 antibody (red) in cerebral cortex (Ctx) and hippocampus (Hip); 4,6-diamidino-2-phenylindole (DAPI) (blue) was used to counterstain cell nuclei. Arrows indicate Fpn1 expression and localization. (D) Immunofluorescence of neuronal and astrocytic cells using anti-GLUT1 (green), anti-GLAST (red) and anti-Fpn1 antibodies in cerebral cortex (Ctx) and hippocampus (Hip). Scale bars: 40X. *statistically significant vs WT A control group *P <0.05; **P<0.01 ***P <0.001 using OneWay ANOVA followed by Bonferroni’s post hoc analysis.

To further investigate this Fpn1 reversal distribution and to better understand Hepc/Fpn1 correlation at the cellular level, we co-labelled Fpn1 with a specific astrocytic and neuronal marker: GLAST and VGLUT1 respectively. We found that Fpn1 increases with aging and co-localizes with astrocytes in the Ctx and Hip, while it remains constant in neurons (Fig 3D); the same co-localization is present also in WT A mice (Fig 1S). These results demonstrate that neurons and astrocytes respond differently to iron excess.

### Ferritin response to aging, iron increase and NCOA4-mediated modification of ferritin heteropolymers

To investigate how neuronal cells responded to the increase of iron amount, we analysed the iron deposit protein Ft in the total brain evaluating separately the two polymers Ft-L and Ft-H. As expected, we observed a significant increase in Ft-L amount (Fig. 4A), but a 40% reduction of Ft-H polymers in WT O animals’ brains compared to the WT A ones (Fig. 4B).

**Figure.4.**
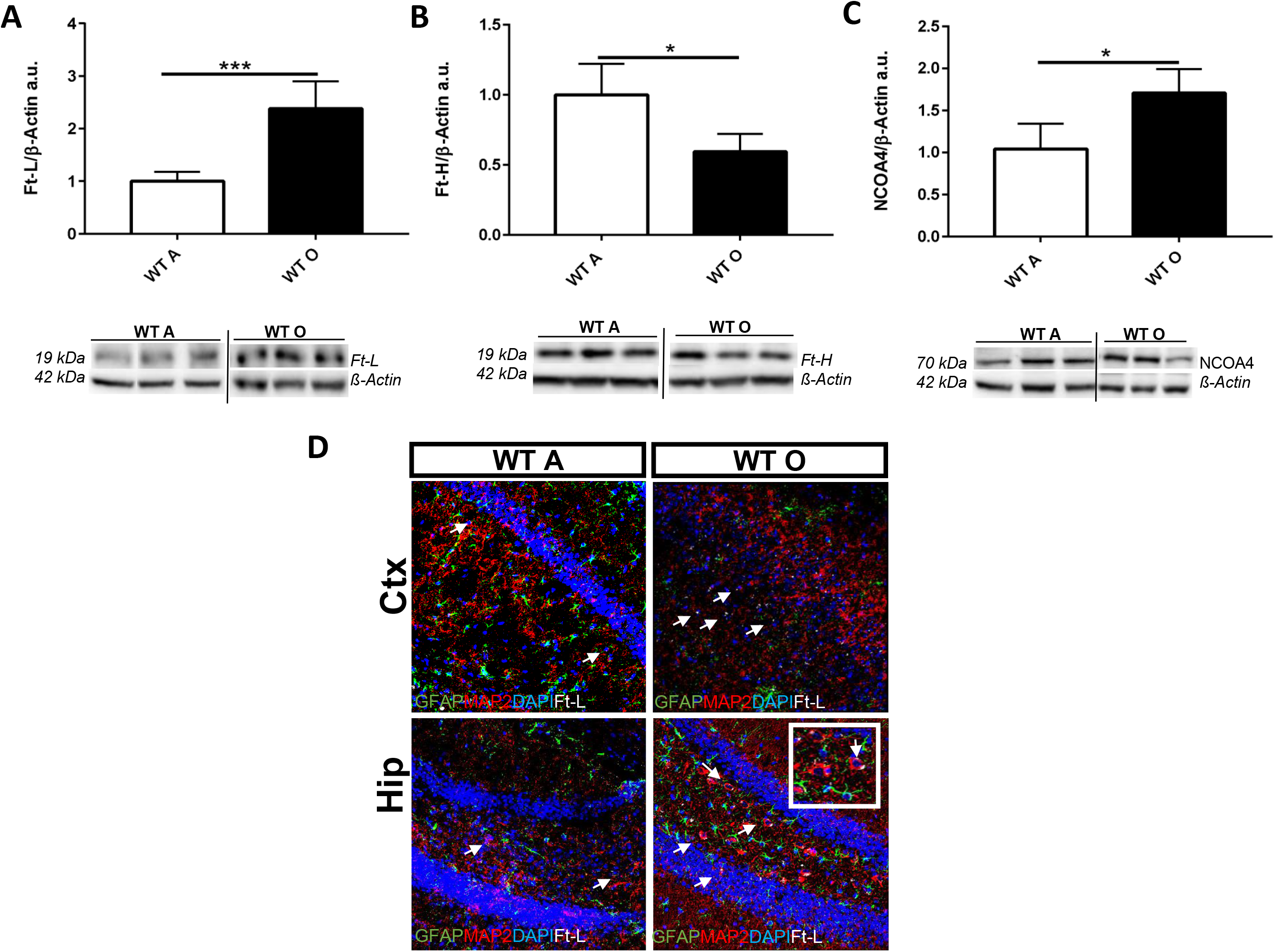
Ferritin’s response to ageing iron increase and NCOA4 involvement. (A) Western blotting analysis and quantification of Ferritin-L (Ft-L), (B) Ferritin-H (Ft-H), and (C) Nuclear receptor coactivator 4 (NCOA4). (D) Immunofluorescence of astrocytic and neuronal cells using anti-GFAP (green), anti-MAP2 (red), and anti-Ft-L antibodies in cerebral cortex (Ctx) and hippocampus (Hip). 4,6-diamidino-2-phenylindole (DAPI) (blue) was used to counterstain cell nuclei. Scale bars: 40X. * statistically significant vs WT A control group *P <0.05; **P <0.01 ***P <0.001 using unpaired T-test.

To verify if these differences were caused by the action of the ferritinophagy inductor NCOA4, we investigated NCOA4 transcription and translation in the brain. Indeed, NCOA4 gene results to be highly transcribed in the brain and its expression level is comparable to that of the liver (Ct values 25±1 and 24±1.5 respectively) (Bellelli 2016 and Fig 2S). More importantly, NCOA4 protein amount is significantly increased in WT O mice brain compared to WT A one (Fig. 4C). To discriminate if the accumulation of iron specifically occurred inside neurons and/or astrocytes, we co-stained Ft-L with MAP-2 and GFAP, respectively. Compared to WT A mice, in WT O mice brains’ sections we observed a specific increase of Ft-L deposits in cortical and hippocampal neurons compared to astrocytes (Fig. 4D).

To date, no data concerning NCOA4 expression levels, function and role in the brain during aging were available. Our results show for the first time that NCOA4 has a considerable transcription in the CNS and that brain cells respond to higher iron amounts increasing NCOA4 production. We hypothesize that NCOA4, acting specifically on Ft-H degradation, can promote the formation of Ft-L rich heteropolymers, more suitable to iron storage.

### Blood Brain Barrier (BBB) permeability increases during animal aging

To verify if the higher iron flux in the CNS is due to an increase of BBB permeability, we analysed ZO-1 protein, a marker of BBB integrity. The role of ZOs proteins is to maintain the compactness of BBB since they act as a bridge connecting the cytoplasmic tails of Claudin and Occludin to the actin cytoskeleton in order to stabilize the tight junction (TJ) structure (Dobrogowska 2004; Maiuolo 2018). We found that the ZO-1 levels of expression significantly decrease during aging (Fig. 5A). We can argue that BBB altered permeability, due to physiological aging, could be responsible for the increase of iron flux from systemic circulation to the brain; this could in turn lead to iron accumulation.

**Figure.5.**
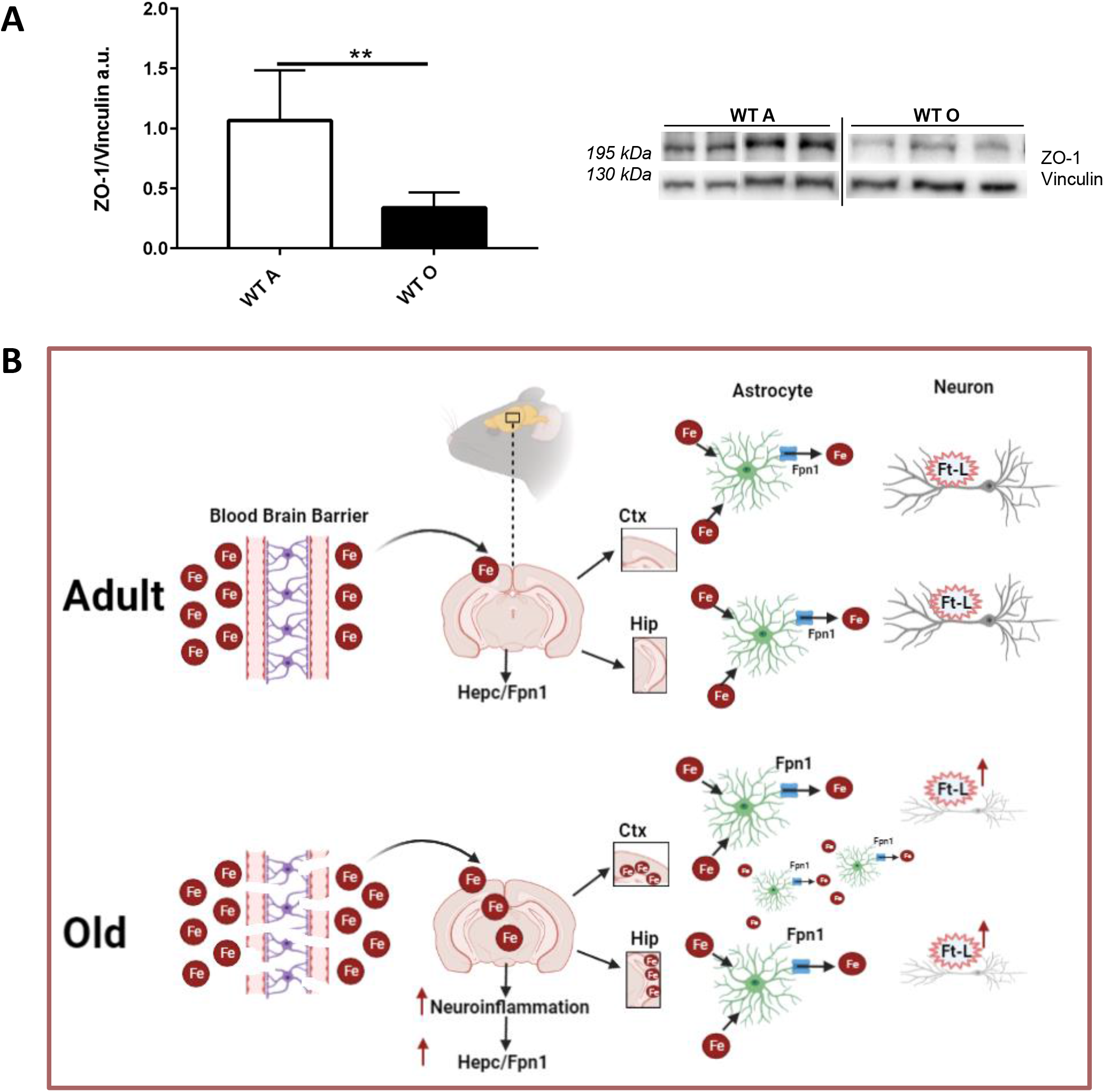
Blood brain barrier (BBB) altered permeability. (A) Western blotting analysis and quantification of Zonula occludens-1 (ZO-1). * statistically significant vs WT A control group (N=3 at least) *P <0.05; **P <0.01 ***P <0.001 using unpaired T-test. (B) Schematic model illustrating iron metabolism in old mice brains vs adults (see text for details). Fe: iron; Hepc: Hepcidin; Fpn 1: Ferroportin 1; Ft-L: Ferritin-L; Ctx: cerebral cortex; Hip: hippocampus.

## Discussion

The CNS exhibits peculiar characteristics in BI management (Zecca 2004). Although iron content should remain roughly stable in the brain during life, an increase in iron content was observed in multiple brain regions during aging (Zecca 2004; Lozoff 2006) and in neurodegenerative diseases with old age onset such as PD and AD (Rouault 2013). In particular, iron overload in motor area, such as basal ganglia, may explain the motor deterioration (Mill 2010) and it may trigger the aggregation of proteins such as α-synuclein (Yamamoto 2002) and the formation of inclusion bodies containing damaged or aggregated proteins that could cause endoplasmic reticulum stress (Liu 2012). Furthermore, in glial cells, the main sensors of neuronal homeostasis alteration, it is iron accumulation which triggers the release of pro-inflammatory cytokines, determining a pro-inflammatory environment which promotes the neurodegenerative process (Ndaysaba 2019).

The major protective barrier against iron overload in the brain is the Blood Brain Barrier (BBB). Indeed, the BBB mitigates iron entry from the blood through highly regulated and selective transport systems; iron is acquired by neurons and glial cells through iron transporter proteins, and released from these cells through the iron export protein Ferroportin 1 (Fpn1) (Rouault 2013). Data show that iron crosses the BBB, which is formed by cerebrovascular endothelial cells (or brain vascular endothelial cells, BVECs), bond to Tf through TfR-mediated endocytosis (Moos 2004; Leitner 2012). BVECs export intracellular iron using Fpn1, whose export activity is conditioned by the action of the iron ferroxidases ceruloplasmin and hephaestin (Rouault 2013).

It is now clear that systemic iron regulation is based on a complex protein regulatory system focused on the hepatic Hepcidin (Hepc) production under different stimuli, such as the low or the high iron availability, hypoxia or inflammation, and on its blocking action on the exporter Fpn1 (Muckenthaler 2017). This complex regulatory system is active also in the CNS (Rouault 2013). Data show that Hepc is expressed in different brain regions, located in both glial cells and neurons, and under BI overload condition, an increase of Hepc corresponds to a Fpn1 decrease (Wang 2010; Ding 2011; Du 2015). However, it is not clear yet whether the source of Hepc in the brain is only local or if hepatic Hepc can pass the BBB and regulate CNS iron availability as well (Vela 2018). Moreover, how does this regulatory system respond to intracerebral iron increase during physiological aging process? To answer to this question, we studied the expression of some proteins that play a central role in maintaining systemic iron homeostasis in C57BL/6 mice brain at different animal ages.

We demonstrated that the progressive increase in iron content that we observed in animals’ brain during aging triggers the onset of an neuroinflammatory condition, measured by an increased expression of SAA-1 (Serum Amyloid protein A1), a protein expressed in acute inflammatory phase and a reliable marker of neuroinflammation (Yu 2019). During acute neuroinflammation, in fact, SAA-1 increases up to 1000 times and can stimulate the release of cytokines and chemokines (Jang 2019). Interestingly, the same protein has been found to be overexpressed in the brain of AD patients, as well as in the AD mouse model APP/SAA, that recapitulates the disease (Jang 2019). Moreover, the increase in transcription of Nrf2, a redox-sensitive transcription factor whose activation results in cellular antioxidant responses via modulation of several stress-responsive proteins (Jiang 2019), supports the evidence of an oxidative stressful condition in our WT O mice brain.

It is known that astrocytes are less susceptible than neurons to the toxicity stimulated by iron accumulation (Kress 2002); indeed, they play a protective role towards neurons in thiamine deficiency-induced neurodegeneration (Ke 2004) during the early stage. Conversely, in the late stage of neurodegeneration, consistent astrocytes activation is visible, demonstrated by the progressive increase of GFAP, together with increased neuroinflammation and neurodegeneration (Ke 2004). Indeed, in line with these findings, in both iron overloaded parenchymal regions of our WT O mice, Ctx and Hip, we observed high astrocytic activation and microglial expression, as a clear sign of neuroinflammation leading to progressive neurodegeneration.

Moreover, as a response to the increased iron amount and/or neuroinflammation in WT M-A and WT O mice, we found an increase of 8-10 fold of Hepc transcription, while Fpn1 amount decreases, even if more gradually compared to Hepc in the whole brain. However, when we restrain the observation to the areas in which iron overload is mainly localized, we found that Fpn1 is increased specifically in WT O mice Ctx and Hip. We can hypothesize that this could be related to the rise of Hepc levels in the other brain regions in which there is no iron overload.

Furthermore, we showed that during animals aging Fpn1 amount in the brain is cell-specific. Indeed, Fpn1 protein increases during aging in cortical and hippocampal astrocytes, but not in neurons, all this regardless of the relevant Hepc increase (30x) in total brain.

This could be due to: i) a detoxifying mechanism carried out by neurons and astrocytes, aimed to store or remove iron excess respectively; ii) an impairment of the Hepc/Fpn1 physiologic metabolism of cortical and hippocampal neurons and astrocytes due to the selective increase of iron amount in these specific brain regions. It could also be hypothesized that the Fpn1 expressed in these cells is one of the different isoforms already characterized, for example an isoform not responsive to Hepc or modulated through different regulatory mechanisms (Drakesmith 2015).

To evaluate the regulation of the increase of iron content in neuronal tissue, we analyzed the iron deposit protein Ft and a recently identified protein, Nuclear Receptor Coactivator 4 (NCOA4).

NCOA4 is involved in Ft degradation and i) its inactivation in mice causes a hepatic iron overload phenotype (Bellelli 2016); ii) Ft levels are enhanced in cellular models in which NCOA4 is silenced, suggesting that ferritin is constantly degraded by an NCOA4-dependent pathway (Fujimaki 2019); iii) NCOA4 promotes autophagic ferritin degradation through its binding to Ft-H subunit (Dowdle 2014; Mancias 2015). Surprisingly, we found an increased amount of NCOA4 in WT O mice’s brains, contrary to what happens in the liver, where iron overload is associated with NCOA4 silencing (Bellelli 2016). Furthermore, evaluating separately the two ferritin polymers Ft-L and Ft-H, we observed an increase of Ft-L and a decrease of Ft-H during mice aging. The same Ft-L increase was evidenced in cortical and hippocampal neurons, but not in astrocytes, in immunofluorescence analysis. This data further supports the hypothesis that there is a cells specific response to increased iron amount: neurons store the metal in Ft-L rich heteropolymers while astrocytes increase iron export through Fpn1. Also, we hypothesize that brain cells respond to higher intracellular iron amount by increasing NCOA4 that could be responsible for a selective degradation of Ft-H, promoting in turn, the formation of Ft-L rich heteropolymers, which are more effective for iron storage.

Neurodegenerative disorders (NDs) are characterized by the impairment of the BBB integrity and, more precisely, by the alteration of the expression of junction proteins (Luissint 2012). Moreover, the onset of NDs can be promoted by iron accumulation that in turn induces oxidative stress (Skjorringe 2012). To verify if the abnormal flux of iron in the CNS during aging was due to an increase of BBB permeability, we analysed the Zonula occludens-1 protein (ZO-1), a marker of BBB integrity. ZOs are cytoplasmic membrane-associated accessory proteins to maintain the TJ structure and are considered sensitive indicators of normal or altered functional states of BBB (Dobrogowska 2004). The significant decrease in ZO-1 levels that we found in WT O animals supports the hypothesis that iron accumulation in the brain during mice aging is due to a BBB permeability alteration.

As resumed in Fig. 5B, the altered permeability of the BBB observed in our murine model during physiological aging allows increased iron passage from systemic circulation to the brain. Iron overload, selectively localized in the Ctx and Hip, triggers a neuroinflammatory response that in turn activates Hepc/Fpn1 pathway. As a consequence of this, an astrocytic activation and an increased Fpn1 amount occur, this implying an increased iron export from these cells. In order to protect themselves, Ctx and Hip neurons increase the expression of ferritin-L (Ft-L) heteropolymers leading to the production of Ft-L enriched ferritins, more suitable to chelate free iron. This iron availability imbalance could cause oxidative damage, stress and could even cause neurodegeneration.

It is well known that NDs are characterized by inappropriate Hepc production (Qian 2020), so a therapeutic approach aimed at modifying Hepc response could be taken in consideration for treatment. Mini-Hepc and Hepc agonists were already injected into mouse models of transfusion dependent thalassemia (TDT), an iron overload condition characterized by high level of erythroferrone, an Hepc inhibitor (Kautz 2014), and low serum Hepc. These drugs allowed an increase in Hepc levels in the serum that determined, de facto, a decrease in iron concentration in serum, spleen, liver and a reduction of erythroferrone levels (Casu 2020).

Another strategy could be to stimulate or inhibit Hepc production by targeting positive or negative regulators such as Bone Morphogenetic Protein 6 (BMP6), an in vivo inducer of systemic Hepc (Muckenthaler 2017), and Transmembrane Serine Protease 6 (TMPRSS6), a potent Hepc inhibitor (Blanchette 2016). Indeed, stimulators and inhibitors of Hepc were used in cultured cells and animal model also to recover from brain damage (Qian 2020) by blocking inflammatory pathways (Huang 2018) and by reducing oxidative stress and consequently Hepc expression (Zhao 2018).

Even though the dyshomeostasis in iron metabolism in some NDs was a secondary event, the finding that iron-related oxidative damage in AD is an early event in the disease process (Ashraf 2020) suggests that the control of iron levels in the brain remains a worthwhile therapeutic target (Bonda 2011). Some clinical studies on Parkinson’s, Alzheimer’s, Huntington’s and Friedreich’s ataxia patients have already been done to reduce BI amount using iron chelators, although with controversial results (Nunez 2018).

In addition, another protein on which it could be theoretically possible to act is NCOA4, the cargo receptor involved in intracellular iron metabolism. Since a constitutive and systemic NCOA4 depletion showed an iron overload phenotype (Bellelli 2016), we can assert that NCOA4 upregulation could be protective in the context of NDs. Obviously, it is essential to expand the understanding of the detailed role of NCOA4 in the brain through the generation of murine models with targeted deletion of NCOA4 in the brain or using other selective gene silencing techniques (Czarnek 2016). Taken together, these data highlight for the first time the involvement of the Hepc/Fpn1 axis and NCOA4 in BI increase during mice aging as a response to a higher iron flux in CNS consequent to a BBB alteration.

Additional research in animal models of NDs are required to study the response to the increased amount of iron in the brain in order to exploit the results for the prevention and clinical management of patients with these diseases.

## Materials and Methods

### Animals

The C57BL/6 mice (from now on indicated with the acronym WT) used for the study were purchased from the Jackson Laboratory. Both male and female mice and at least three animals for each experimental group were analysed and grouped according to their age. To classify them we used “The Jackson Laboratory” classification (https://www.jax.org). Mice are grouped as follows: until 2 months of age mice are considered Young (WT Y); from 2 to 6 months of age Adult (WT A); from 6 to 12 months of age Middle-aged (WT M-A) and from12 months of age (between 18-24 months of age) Old (WT O) (Fig. 1A). Mice were housed in transparent conventional polycarbonate cages (Tecnoplast, Buggirate, Italy) provided with sawdust bedding, boxes/tunnels hideout as environmental enrichment. Food and water were provided ad libitum; environmental conditions were 12 h/12 h light/dark cycle, room temperature 24 °C ± 1 °C and room humidity 55% ± 5%. Each group of mice was fed with a Standard Diet (SD) (VRF1, Special Diets Services, Essex, United Kingdom). Mice were anaesthetized (ketamine, 100 mg/kg; Ketavet, Bayern, Leverkusen, Germany; xylazine, 5 mg/kg; Rompun; Bayer, Milan, Italy) and sacrificed by cervical dislocation. A subset of 3 WT A and 3 WT O mice were transcardially perfused with 4% paraformaldehyde (PFA) in phosphate buffered saline (PBS). Animals housing and all the experimental procedures were performed in accordance with European (Official Journal of the European Union L276 del 20/10/2010, Vol. 53, p. 33–80) and National Legislation (Gazzetta Ufficiale n° 61 del 14/03/2014, p. 2–68) for the protection of animals used for scientific purposes and the experimental procedure was approved by the Ethical Committee of the University of Turin.

### Real-time quantitative PCR

The brain tissues of the mice were homogenized in TRIzol reagent, separated with chloroform and the total RNA was precipitated with isopropanol. After washing with 70% ethyl alcohol, RNase free water was added to dissolve RNA. For reverse transcription, 2 μg of total RNA, 25 μM random hexamers and 100 U of reverse transcriptase (Applied Biosystems, California, USA) were used. Gene expression levels were measured in the total brain using Real-time quantitative PCR in a CFX96 Real-time System (Bio-Rad, California, USA). For Nuclear factor erythroid 2-related factor 2 (Nfr2) and NCOA4 gene analysis, SYBR Green PCR technology (EVAGreen, Bio-Rad, California, USA) was used with specific primers designed (sequences are reported in Supplemental Material) (Table 1S). For Hepc and Serum Amyloid A1 (SAA1) genes analysis Taqman PCR method was used (Assays-on-Demand, Gene Expression Products, Applied Biosystems, California, USA). In both cases, β-glucuronidase (Gus-β) gene was used as housekeeping control. The results were analyzed using the ΔΔCt method (Livak 2001). Real-time quantitative PCR of all the transcripts were carried out in duplicate.

### Immunoblotting

The Fpn1, Ft-H, Ft-L, NCOA4 and Zonula occludens-1 (ZO-1) proteins amount in the brain was evaluated by Western Blotting (WB) analysis using specific antibodies and normalized to β-Actin protein. 50 μg of total brain lysates were separated on 6–12% SDS polyacrylamide gel and immunoblotted according to standard protocols (Boero 2015). The following antibodies were used to detect the different proteins: Fpn1 (G-16), β-Actin (C-4) and NCOA4 or ARA 70 (H-300) (Santa Cruz Biotechnology, Dallas, Texas, USA); ZO-1 (GeneTex, California, USA). Antibodies used to specifically detect the two ferritin isoforms, Ft-H and Ft-L, were gently provided by Sonia Levi, University of Vita Salute, Milan, Italy. Data from each protein quantification (Image Lab 4.0.1 Software, Bio-Rad, California, USA) were normalized on β-Actin amount in the same samples.

### Immunofluorescence

To perform histological analysis, animals were perfused, brains were removed and post-fixed in PFA for 24h at 4°C, cryoprotected in 30% sucrose in 0.12 M phosphate buffer and processed according to standard protocols. Brains were cut in 30 μm thick coronal sections collected in PBS and then stained to detect the expression of Fpn1 (G-16, Santa Cruz Biotechnology, Dallas, Texas, USA), Ft-L (S. Levi, University of Vita Salute, Milan), Glial Fibrillary Acidic Protein (GFAP) (Dako, California, United States), Microtubule-Associated Protein 2 (MAP-2) (Merck Millipore Burlington, Massachusetts, United States), Vesicular Glutamate Transporter 1 (VGLUT1) (Merck Millipore Burlington, Massachusetts, United States), Glutamate Transporter (GLAST) (Thermo Fischer Scientific Waltham, Massachusetts, United States) and Ionized calcium-binding adaptor molecule 1 (IBA-1) (Abcam, Cambridge, United Kingdom). Incubation with primary antibodies was made overnight at 4°C in PBS with 2% normal donkey serum (NDS) (D’Errico 2013). The sections were then exposed to secondary Cy2-, Cy3-(Jackson ImmunoResearch Laboratories, West Grove, PA) and 647 Alexa Fluor-conjugated antibodies (Molecular Probes Inc, Eugene Oregon) for 1 h at room temperature (RT). DAPI (4,6-diamidino-2-phenylindole, Fluka, Italy) was used to counterstain cell nuclei. After processing, sections were mounted on microscope slides with Tris-glycerol supplemented with 10% Mowiol (Calbiochem, LaJolla, CA). The samples were examined by a Leica TCS SP5 confocal laser scanning microscope (Leica, Mannheim); for each image z-stacks were taken at 40X and 63X magnification.

### Iron parameters

Brain tissues nonheme iron content (BIC) was performed according to standard procedure. Iron amount was evaluated using a colorimetric technique utilizing 20 mg of dissected and dried murine whole brains (Roetto 2010).

For staining of perfused brains nonheme ferrous iron Prussian blue Perl’s staining was performed using a specific commercial kit (Bio-Optica, Milan, Italy). To improve the sensitivity of Perls’ method, an intensification step with DAB (3-3’-diaminobenzidine tetrahydrochloride) (Meguro 2007) was performed.

Images were taken at 10X magnification using a Leica DM4000B automated vertical microscope and the IM50 program for image acquisition (Leica Microsystems, Wetzlar, Germany).

### Statistical analysis

To verify if the differences of mRNA expression and protein production were statistically significant, one-way ANOVA followed by Bonferroni’s post hoc analysis or unpaired T-test were applied according to the experimental group’s number. In both cases, the p values of <0.05 were considered as statistically significant. Analyses were performed with Image Lab 4.0.1 and GraphPad Prism 7.00. Data were expressed as average ± standard error of the mean. WT adult (A) mice were used as normalizer.

## Author Contributions

MM performed research, analyzed data and wrote the paper. GA* and MB performed research and analyzed data. SS and AR designed and performed research, analyzed data and wrote the paper. All the authors contributed to the article and approved the submitted version. *Currently PhD student at Department of Molecular Biotechnology and Health Sciences, University of Turin, Italy.

## Acknowledgments

We are grateful to our colleague Sonia Levi for providing us anti Ferritin antibodies.

## Abbreviations

ATP: adenosine triphosphate
ROS: Reactive Oxidative Species
Tf: Transferrin
TfR1: Transferrin Receptor 1
DMT1: Divalent Metal Transporter 1
Fpn1: Ferroportin 1
Ft-H: Ferritin heavy chain
Ft-L: Ferritin light chain
Hep: Hepcidin
BBB: Blood Brain Barrier
Tf-Fe2: Transferrin iron complex
BMVECs: Brain microvascular endothelial cells
CNS: Central Nervous System
IRPs: Iron Regulatory Proteins
NEO-1: Neogenin-1
BI: Brain Iron
miRNA: MicroRNAs
NDs: Neurodegenerative disorders
PD: Parkinson’s disease
AD: Alzheimer disease
MS: Multiple sclerosis
NCOA4 or ARA70: Nuclear Receptor Coactivator 4 or Androgen Receptor (AR) Coactivator
HERC2: E3 ubiquitin protein ligase
SD: Standard Diet
Nfr2: Nuclear factor Erythroid 2-related factor 2
SAA1: Serum Amyloid A1
Gus-β: β-glucuronidase
ZO-1: Zonula Occludens-1
GFAP: Glial Fibrillary Acidic Protein
MAP-2: Microtubule-Associated Protein 2
VGLUT1: Vesicular Glutamate Transporter 1
GLAST: Glutamate Transporter
IBA-1: Ionized calcium-binding adaptor molecule 1
NDS: normal donkey serum
DAPI: 4,6-diamidino-2-phenylindole
BIC: Brain iron content
DAB: 3-3-diaminobenzidine tetrahydrochloride
Ctx: Cerebral cortex
Hip: Hippocampus
3V: third ventricle
WT Y: wild type Young
WT A: wild type Adult
WT M-A: wild type Middle-aged
WT O: wild type Old
ALS: Amyotrophic Lateral Sclerosis
TJ: Tight junction
BVECs: brain vascular endothelia cells
TDT: transfusion dependent thalassemia
BMP6: Bone Morphogenetic Protein 6
TMPRSS6: Transmembrane Serine Protease 6

## References

Almutairi M, Chen G, Yuexian G X, Yanzhong C, Honglian S (2016) Factors controlling permeabilityof the Blood-Brain Barrier. Cell Mol Life Sci 73: 57–77

Ashraf A, Jeandriens J,Parkes HG, So PW (2020) Iron dyshomeostasis, lipid peroxidation and perturbedexpression of cystine/glutamate antiporter in Alzheimer’s disease: Evidence of ferroptosis. Redox Biol32: 101494

Arosio P, Ingrassia R, Cavadini P (2009) Ferritins: a family of molecules for iron storage, antioxidationand more. Biochim Biophys Acta 1790: 589–99

Bellelli R, Federico G, Matte A, Colecchia D, Iolascon A, Chiariello M, Santoro M, De Franceschi L,Carlomagno F (2016) NCOA4 Deficiency Impairs Systemic Iron Homeostasis Cell Rep 26;14: 411–421.

Biasiotto G, Di Lorenzo D, Archetti S, Zanella (2016) I Iron and Neurodegeneration: Is Ferritinophagythe Link? Mol Neurobiol 53: 5542–74

Bien-Ly N, Yu YJ, Bumbaca D, Elstrott J, Boswell CA, Zhang, Y, Luk W, Lu Y, Dennis MS et al (2014) Transferrin receptor (TfR) trafficking determines brain uptake of TfR antibody affinity variants J ExpMed 10;211: 233–44

Blanchette NL, Manz DH, Torti FM, Torti SV (2016) Modulation of Hepc to treat iron deregulation: potential clinical applications. Expert Rev Hematol 9: 169–86

Block ML, Hong JS (2005) Microglia and inflammation-mediated neurodegeneration: multiple triggerswith a common mechanism. Prog Neurobiol 76: 77–98

Boero M, Pagliaro P, Tullio F, Pellegrino RM, Palmieri A, Ferbo L, Saglio G, De Gobbi M, Penna C, Roetto A (2015) A comparative study of myocardial molecular phenotypes of two tfr2β null mice: role in ischemia/reperfusion. Biofactors 41: 360–71

Bonda DJ, Lee HG, Blair JA, Zhu X, Perry G, Smith MA (2011) Role of metal dyshomeostasis in Alzheimer’s disease. Metallomics 3: 267–270

Buijs M, Doan NT, Van Rooden S, Versluis MJ, Lew BV, Milles J, Grond JVD, Buchem MAV (2017)In vivo assessment of iron content of the cerebral cortex in healthy aging using 7-Tesla T2*-weightedphase imaging. Neurobiol Aging 53: 20–26

Bulet P, Stöcklin R, Menin L (2004) Anti-microbial peptides: from invertebrates to vertebrates. ImmunolRev 198: 169–84

Casu C, Chessa R, Liu A, Gupta R, Drakesmith H, Fleming R, Ginzburg YZ, MacDonald B, Rivella S (2020) MiniHepcs improve ineffective erythropoiesis and splenomegaly in a new mouse model of adult ß-thalassemia major. Haematologica 105: 1835–1844

Conde JR, Streit WJ (2006) Microglia in the aging brain. J Neuropathol Exp Neurol 65: 199–203

Czarnek M, Bereta J (2016) The CRISPR-Cas system - from bacterial immunity to genome engineering. Postepy Hig Med Dosw (Online)70: 901–16

D’Errico P, Boido M, Piras A, Valsecchi V, De Amicis E, Locatelli D, Capra S, Vagni F, Vercelli A, Battaglia G (2013) Selective vulnerability of spinal and cortical motor neuron subpopulations in delta7 SMA mice. PLoS One 8: e82654

Ding H, Yan CZ, Shi H, Zhao YS, Chang SY, Yu P, Wu WS, Zhao CY, Chang YZ, Duan LX (2011)Hepc is involved in iron regulation in the ischemic brain. PLoS One 6: e25324

Dobrogowska DH, Vorbrodt AW (2004) Immunogold localization of tight junctional proteins in normaland osmotically-affected rat blood-brain barrier. J Mol Histol 35: 529–39

Donovan A, Brownlie A, Zhou Y, Shepard J, Pratt SJ, Moynihan J, Paw BH, Drejer A, Barut B, ZapataA et al (2000). Positional cloning of zebrafish ferroportin1 identifies a conserved Vertebrate iron exporter. Nature 17;403: 776–81

Dowdle WE, Nyfeler B, Nagel J, Elling RA, Liu S, Triantafellow E, Menon S, Wang Z, Honda A, Pardee G et al (2014) Selective VPS34 inhibitor blocks autophagy and uncovers a role for NCOA4 in ferritin degradation and iron homeostasis in vivo. Nat Cell Biol 16: 1069–79

Drakesmith H, Nemeth E, and Ganz T (2015) Ironing out ferroportin. Cell Metab 3;22: 777–87

Du F, Qian ZM, Luo Q, Yung WH, Ke Y (2015) Hepc Suppresses Brain Iron Accumulation by Downregulating Iron Transport Proteins in Iron-Overloaded Rats. Mol Neurobiol 52: 101–14

Farrall AJ, Wardlaw JM (2009) Blood-brain barrier: aging and microvascular disease--systematic reviewand meta-analysis. Neurobiol Aging 30: 337–52

Fujimaki M, Furuya N, Saiki S, Amo T, Imamichi Y, Hattori N (2019) Iron Supply via NCOA4-Mediated Ferritin Degradation Maintains Mitochondrial Functions. Mol Cell Biol 27;39: 00010–19

Ganz, T (2013) Systemic iron homeostasis. Physiol Rev 93: 1721–41

Ganz T, Nemeth E (2012) Iron homeostasis and its disorders in mice and men: potential lessons for rhinos. J Zoo Wildl Med 4: S19–26

Ginzburg YZ (2019) Hepc-ferroportin axis in health and disease. Vitam Horm 110: 17–45

Glass CK, Saijo K, Winner B, Marchetto MC, Gage FH (2010) Mechanisms underlying inflammation in neurodegeneration. Cell 140: 918–34

Hirsch EC, Hunot S. Neuroinflammation in Parkinson’s disease: a target for neuroprotection? (2009) Lancet Neurol 8:382–97

Huang SN, Ruan HZ, Chen, MYJ, Zhou G, Qian ZM (2018) Aspirin increases ferroportin 1 expressionby inhibiting Hepc via the JAK/STAT3 pathway in interleukin 6-treated PC-12 cells. Neurosci Lett 662: 1–5

Jang S, Young Jang W, Choi M, Lee J, Kwon W, Yi J, Park S, Yoon D, Lee S, Kim MO et al (2019) Serum amyloid A1 is involved in amyloid plaque aggregation and memory decline in amyloid beta abundant condition. Transgenic Res 28: 499–508

Jiang Z, Wang J, Liu C, Wang X, Pan J (2019) Hyperoside alleviated. N-acetyl-para-amino-phenol-induced acute hepatic injury via Nrf2 activation. Int J Clin Exp Pathol 12: 64–76

Kautz L, Jung G, Valore E, Rivella S, Nemeth E, Ganz T (2014) Identification of Erythroferrone as an Erythroid Regulator of Iron Metabolism. Nat Genet 46: 678–84

Ke ZJ, Gibson GE (2004) Selective response of various brain cell types during neurodegeneration induced by mild impairment of oxidative metabolism. Neurochem Int 45: 361–9

Kress GJ, Dineley KE, Reynolds IJ (2002) The relationship between intracellular free iron and cell injury in cultured neurons, astrocytes, and oligodendrocytes. J Neurosci 22: 5848–55

Leibold EA, Gahring LC, Rogers SW (2001) Immunolocalization of iron regulatory protein expressionin the murine central nervous system. Histochem Cell Biol 115: 195–203

Leitner DF, Connor JR (2012) Functional roles of transferrin in the brain. Biochim Biophys Acta 1820:393–402

Lier J, Winter K, Bleher J, Grammig J, Mueller WC, Streit W, Bechmann I (2019) Loss of IBA1-Expression in brains from individuals with obesity and hepatic dysfunction. Brain Res 1710: 220–229

Liu Y, Connor JR (2012) Iron and ER stress in neurodegenerative disease. Biometals 25: 837–45

Livak KJ, Schmittgen TD (2001) Analysis of relative gene expression data using real-time quantitativePCR and the 2(-Delta Delta C(T)) Method. Methods 25: 402–8

Lozoff B, Georgieff MK (2006) Iron deficiency and brain development. Semin Pediatr Neurol 13: 158–165

Luissint AC, Artus C, Glacial F, Ganeshamoorthy K, Couraud PO (2012) Tight junctions at the blood brain barrier:physiological architecture and disease-associated dysregulation. Fluids Barriers CNS 9;9:23

Maiuolo J, Micaela G, Vincenzo M, Scicchitano M, Carresi C, Scarano F, Bosco F, Nucera S, Ruga S, Zito MC et al (2018) The “Frail” Brain Blood Barrier in Neurodegenerative Diseases: Role of EarlyDisruption of Endothelial Cell-to-Cell Connections Int J Mol Sci 10;19:2693

Mancias JD, Pontano Vaites LP, Nissim S, Biancur DE, Kim AJ, Wang X, Liu Y, GoesslingW, Kimmelman AC, Harper JW et al (2015) Ferritinophagy via NCOA4 is required for erythropoiesis and is regulated by iron dependent HERC2-mediated proteolysis. Elife 5;4:e10308

Mancias JD, Wang X, Gygi SP, Harper JW, Kimmelman AC (2014) Quantitative proteomics identifiesNCOA4 as the cargo receptor mediating ferritinophagy. Nature 509: 105–9

McCarthy RC, Kosman DJ (2014) Glial cell ceruloplasmin and Hepc differentially regulate iron effluxfrom brain microvascular endothelial cells. PLoS One 12;9:e89003

Meguro R, Asano Y, Odagiri S, Li C, Iwatsuki H, Shoumura K (2007) Nonheme-iron histochemistry forlight and electron microscopy: a historical, theoretical and technical review. Arch Histol Cytol 70: 1–19

Mills E, Dong XP, Wang F, Xu H (2010) Mechanisms of BI transport: insight into neurodegeneration and CNS disorders. Future Med Chem 2: 51–64

Morello N, Tonoli E, Logrand F, Fiorito V, Fagoonee S, Turco E, Silengo L, Vercelli A, Altruda F, Tolosano E (2009) Haemopexin affects iron distribution and ferritin expression in mouse brain. J Cell Mol Med 13: 4192–204

Moos T (1996) Immunohistochemical localization of intraneuronal transferrin receptor immunoreactivity in the adult mouse central nervous system. J Comp Neurol 375: 675–92

Moos T, Morgan EH (2004) The metabolism of neuronal iron and its pathogenic role in neurological disease. Ann N Y Acad Sci 1012: 14–26.

Moos T, Rosengren Nielsen T, Skjørringe T, Morgan EH (2007) Iron trafficking inside the brain. J Neurochem 103: 1730–40

Mosley RL, Benner EJ, Kadiu I, Thomas M, Boska MD, Hasan K, Laurie C, Gendelman HE (2006) Neuroinflammation, Oxidative Stress and the Pathogenesis of Parkinson’s Disease. Clin Neurosci Res 6: 261–81

Muckenthaler U, Rivella S, Hentze W, Galy B (2017) A Red Carpet for Iron Metabolism. Cell 168: 344–361

Nai A, Lidonnici MR, Federico G, Pettinato M, Olivari V, Carrillo F, Crich SG, Ferrari G, CamaschellaC, Silvestri L et al (2021) NCOA4-mediated ferritinophagy in macrophages is crucial to sustain erythropoiesis in mice. Haematologica 106: 795–805

Ndaysaba A, Kaindlstorfer C, and Wenning G (2019) Iron in neurodegeneration-Cause or Consequence? Front Neurosci 1;13:180

Nunez M, Chana-Cuevas P (2018) New Perspectives in Iron Chelation Therapy for the Treatment of Neurodegenerative Diseases. Pharmaceuticals (Basel) 19;11:109

O’Callaghan JP, Sriram K (2005) Glial fibrillary acidic protein and related glial proteins as biomarkers of neurotoxicity. Expert Opin Drug Saf 4: 433–42

Pellegrino RM, Boda E, Montarolo F, Boero M, Mezzanotte M, Saglio G, Buffo A, Roetto A (2016) Transferrin Receptor 2 Dependent Alterations of BI Metabolism Affect Anxiety Circuits in the Mouse. Sci Rep 1;6:30725.

Qian ZM, Ke Y (2020) Hepc and its therapeutic potential in neurodegenerative disorders. Med Res Rev 40: 633–653

Quiles Del Rey M, Mancias JD (2019) NCOA4-Mediated Ferritinophagy: A Potential Link to Neurodegeneration. Front Neurosci 14;13:238

Raha-Chowdhury R, Raha AA, Forostyak S, Zhao JW, Stott SR, Bomford A (2015) Expression and cellular localization of Hepc mRNA and protein in normal rat brain. BMC Neurosci 21;16:24

Rodriguez A, Pan P, Parkkila S (2007) Expression studies of neogenin and its ligand hemojuvelin in mouse tissues. J Histochem Cytochem 55: 85–96

Roetto A, Di Cunto F, Pellegrino RM, Hirsch E, Azzolino O, Bondi A, Defilippi I, Carturan S, Miniscalco B, Riondato F et al (2010) Comparison of 3 Tfr2-deficient murine models suggests distinct functions for Tfr2-alpha and Tfr2-beta isoforms in different tissues. Blood 115: 3382–9

Roetto A, Mezzanotte M, Pellegrino RM (2018) The Functional Versatility of Transferrin Receptor 2 and Its Therapeutic Value. Pharmaceuticals (Basel) 11:115

Rouault TA (2013) Iron metabolism in the CNS: implications for neurodegenerative diseases. Nat Rev Neurosci 14: 551–64

Sangkhae V, Nemeth E (2017) Regulation of the Iron Homeostatic Hormone Hepc. Adv Nutr 8: 126–136

Santana-Codina N, Mancias JD (2018) The Role of NCOA4-Mediated Ferritinophagy in Health and Disease. Pharmaceuticals (Basel) 23;11:114

Skjorringe T, Moller LB, Moos T (2012) Impairment of interrelated iron and copper homeostatic mechanisms in brain contributes to the pathogenesis of neurodegenerative disorders. Front Pharmacol 25;3:169

Thirupathi A, Chang YZ (2019) Brain Iron Metabolism and CNS Diseases. Adv Exp Med Biol 1173: 1–19

Vela D (2018) Hepc, an emerging and important player in Brain Iron homeostasis. J Transl Med 7;16:25

Vezzani A, Aronica E, Mazarati A, Pittman QJ (2013) Epilepsy and brain inflammation. Exp Neurol. 244: 11–21

Wang SM, Fu LJ, Duan XL, Crooks DR, Yu P, Qian ZM, Di XJ, Li J, Rouault TA, Chang YZ (2010) Role of Hepc in murine BI metabolism. Cell Mol Life Sci 67:123–33

Ward R, Zucca F, Duyn J, Crichton R, Zecca L (2010) The role of iron in brain aging and neurodegenerative disorders. Lancet Neurol 13: 1045–60

Yamamoto A, Shin RW, Hasegawa K, Naiki H, Sato H, Yoshimasu F, Kitamoto T (2002) Iron (III) induces aggregation of hyperphosphorylated tau and its reduction to iron (II) reverses the aggregation: implications in the formation of neurofibrillary tangles of Alzheimer’s disease. J Neurochem 82: 1137–47

Yu M, Li Q, Mo S, Ni Y, Han F, Wang YB, Tu YX (2019) SAA1 increases NOX4/ROS production to promote LPS-induced inflammation in vascular smooth muscle cells through activating p38MAPK/NF-κB pathway. BMC Mol Cell Biol 19; 20:15

Zecca L, Youdim MBH, Riederer P, Connor JR, Crichton RR (2004) Iron, brain aging and neurodegenerative disorders. Nat Rev Neurosci 5: 863–73

Zechel S, Huber-Wittmer K, von Bohlen und Halbach O (2006) Distribution of the iron-regulating protein Hepc in the murine central nervous system. J Neurosci Res 84: 790–800

Zhao Y, Xin Z, Li N, Chang S, Chen Y, Geng L, Chang H, Shi H, Chang YZ (2018) Nano-liposomes oflycopene reduces ischemic brain damage in rodents by regulating iron metabolism. Free Radic Biol Med 124: 1–11

